# BAX is necessary for neuronal death following exposure to isoflurane during the neonatal period

**DOI:** 10.1101/2020.08.25.267120

**Authors:** Andrew M. Slupe, Laura Villasana, Kevin M. Wright

**Affiliations:** Department of Anesthesiology and Perioperative Medicine, Oregon Health & Science University, Portland, OR, United States of America; Vollum Institute, Oregon Health & Science University, Portland, OR, United States of America

**Author notes:** Author Contributions: AMS and KMW participated in overall study design and execution as well as manuscript preparation. LV participated in animal behavioral experiments and manuscript preparation.

## Abstract

Exposure to volatile anesthetics during the neonatal period results in acute neuronal death in rodent and non-human primate models, potentially leading to lasting cognitive deficits. We used *Bax*^*-/-*^ mice to show that neuronal death following neonatal exposure to isoflurane is mediated by the apoptotic pathway, and that GABAergic interneurons are selectively vulnerable. Neonatal *Bax*^*-/-*^ mice also showed attenuated microglial activation after exposure to isoflurane, indicating that neuroinflammatory response is secondary to neuronal apoptosis. Isoflurane-induced neuronal apoptosis in neonates appeared to have little effect on seizure threshold or cognitive function later in life. Collectively, these findings define the acute injury mechanism of volatile anesthetics during the neonatal period.

## Introduction

Exposure to volatile anesthetics in early life has consistently been found to result in widespread neuronal death in small mammal and non-human primate models(1). Vulnerability to anesthesia-associated cell death is confined to a temporal window coincident with high levels of brain growth and synaptogenesis(2). Human correlates of these processes suggest that vulnerability to injury for pediatric patients receiving anesthesia may extend into early childhood(3, 4). Neuronal injury by volatile anesthetics may be responsible for lasting behavioral and learning deficits in children exposed to the perioperative environment(5). The molecular pathways involved in neuronal death following exposure to volatile anesthetics in early life remain unclear.

Several prior observations have suggested that apoptosis may be the dominant mechanism of neuronal death following exposure to volatile anesthetics in the neonatal period. Anesthesia exposure alters the expression ratio of Bcl-2 family members in favor apoptosis through down-regulation of anti-apoptotic Bcl-2 and Bcl-x(L) and up regulation of pro-apoptotic Bax and Bad(2, 6-8). The morphology of affected neurons is consistent with apoptotic cell death(9). Finally, early life exposure to ethanol, which has a mechanism of action believed to be similar to volatile anesthetics, results in neuroapoptosis(10). These studies provide circumstantial evidence, but do not definitively demonstrate, that neuronal death following exposure to volatile anesthetics in early life occurs by apoptosis.

Vulnerability to anesthesia-associated death may not be uniform across all neuronal types. It has been suggested that GABAergic interneurons in superficial cortical layers may be overrepresented among the dying neurons(9). GABAergic interneurons normally undergo significant cellular pruning by apoptosis during the same developmental window as vulnerability to anesthesia(16). Therefore, increased susceptibility to anesthesia may reflect an exacerbation of a normal physiological process. It is unknown how exposure to anesthesia impacts this normal physiological cell loss and what the consequences of neuron loss may be on neuronal circuit function.

Bax is activated by signaling events in cells undergoing apoptotic cell death(11). In neurons, genetic deletion of *Bax* is sufficient to block the apoptotic pathway in response to multiple stimuli, including trophic factor withdrawal, excitotoxic insult, and ethanol exposure(10, 12, 13). Blocking Bax activation is therefore a putative target to delineate neuronal death occurring through activation of the apoptotic pathway from other mechanisms of injury. Following early life exposure to ethanol, a neuroinflammatory response characterized by microglial activation occurs as a result of Bax-mediated apoptosis(14). A similar inflammatory profile following exposure to volatile anesthetics has been observed(15). However, it is unknown if the neuroinflammatory response to volatile anesthetics occurs as a consequence of neuronal death, or if it is an independent process that contributes to neuronal death.

In this study, we examined the mechanism of neurotoxicity and neuroinflammation following exposure to isoflurane in early life. We show that neuronal apoptosis following exposure to isoflurane requires Bax function, and that microglial activation is secondary to neuronal apoptosis. We profiled GABAergic neuron vulnerability to anesthesia-induced death, and probed for a disruption of global inhibitory-excitatory balance later in life. Finally, we sought to determine whether neuronal loss due to neonatal exposure to anesthesia affected inhibitory-excitatory balance or cognitive function later in life.

## Materials and Methods

### Animals and anesthetic exposure

All procedures involving animals were approved by the Oregon Health & Science University Institutional Animal Care and Use Committee and conformed to the National Institutes of Health’s *Guide for the Care and Use of Laboratory Animals*. Animals were allowed *ad lib* access to food and water and maintained in standard 12-hour light-dark cycle. Heterozygous *Bax* mice (B6.129×1-Bax^tm1Sjk^lj, stock #002994), *Bax*^*Flox*^;*Bak*^*-/-*^ (B6;129-Baxtm2Sjk Bak1tm1Thsn/J, stock #006329), *Rosa26*^*LSL-TdTomato*^ (B6.Cg-Gt(ROSA)26Sortm9(CAG-tdTomato)Hze/J, stock #007909), and *Gad2-IRES-Cre* (Gad2tm2(cre)Zjh/J, stock #010802) were obtained from The Jackson Laboratory. A conditional GABAergic interneuron specific *Bax* knock-out/reporter line, *Gad2*^*IRES- Cre*^;*Bax*^*Flox*^; *Rosa26*^*TdTom/TdTom*^, was generated by interbreeding of *Bax*^*Flox*^;*Bak*^*-/-*^, *Rosa26*^*LSL-TdTomato*^ and *Gad2-IRES-Cre*. This breeding strategy restored the *Bak* allele, so all mice were either *Bak*^*+/+*^ or *Bak*^*+/-*^. Heterozygosity of the *Gad2* allele was preserved in the experimental population, as altered seizure threshold has been reported in homozygous *Gad2-IRES-Cre* animals(17). Proper recombination by the *Gad2-IRES-Cre* line was verified by expression of tdTomato specifically in GABAergic populations in all experimental animals.

On PND 7, neonatal mice were exposed to isoflurane partially titrated to a level of 1 MAC for 6 hours. At the beginning of the 12-hour light cycle animals were placed on soft bedding in an acrylic induction chamber on circulating water bath heaters such that chamber temperature was maintained at 34°C. Humidity within the chamber was maintained with an open water bath placed near the common gas inlet. Carrier gas was composed of medical air and oxygen at an FiO_2_ of 50% verified with a MaxO_2_ ME oxygen sensor (MaxTec, Salt Lake City, UT) delivered at 1 LPM. Isoflurane concentration within the induction chamber was monitored using a POET II gas analyzer (Criticare Technologies, Inc, North Kingsworth, RI) and sampling from the chamber exhaust port. Our experience and previous reports suggest that the potency of isoflurane increases with prolonged exposure in rodents, with resultant high levels of mortality during exposure without titration(18). For this reason the exposure paradigm used here was partially titrated to a level of 1 MAC guided by previous reports(19). At 150 minutes of exposure responsiveness of the exposed population was assessed by toe pinch and it was found that ∼50% of the exposed animals responded with movement following stimulation. This exposure paradigm was associated with a 1% mortality rate. Control littermates were maintained in similar conditions without isoflurane, all responded to toe pinch at 150 minutes and mortality rate was 0%. Arterial blood gas analysis was not preformed due to institutional prohibitions against un-anesthetized terminal blood sampling. Following completion of the 6 hour exposure isoflurane administration was discontinued, carrier gas flow continued and mice were allowed to recover for 20 minutes following complete washout of isoflurane from the induction chamber. At the end of 20 minutes, responsiveness to toe pinch with vocalization and/or purposeful movement was confirmed and animals were returned to their home cage.

### Genotype and sex determination

DNA was extracted from tissue samples with Extracta DNA Prep kit (QuataBio, Beverly, MA) and PCR reactions were performed using genotype-specific primer sets. Sex was determined using primers directed towards the sex-associated gene *Ube1y1* as previously described(20).

### Neuroapoptosis assessment

Animals were euthanized by rapid decapitation two hours after conclusion of the isoflurane exposure. Brains were isolated and preserved in 4% PFA for 16-24 hours at 4 °C. 50 μm thick coronal sections from were cut on a Leica VT1200 Vibratome. Every fifth section from Paxinos plate P6 #25 to #34 was collected for analysis(21). For immunohistochemistry, free-floating sections were blocked and permeabilized with 2% donkey serum in 0.2% Triton X-100/PBS for four hours at room temperature, followed by antibody staining with 1:1000 anti-cleaved caspase-3 (Cell Signaling Technology) and subsequently 1:1000 anti-rabbit Alexa 568 (ThermoFisher) and 1:5000 Hoechst 33342 (ThermoFisher). Sections were mounted on PermaFrost Plus slides (ThermoFisher), coated in FluoroMount-G (SouthernBiotech) and coverslipped. For Fluorojade C staining, sections were mounted to PermaFrost Plus slides and air dried for 24 hours. If not immediately processed, these slides were stored at −80 °C. Fluorojade C staining was carried out as previously described(22). Following staining sections were coated in DPX Mounting Media (SigmaAldrich) and coverslipped. Sections were imaged with a Zeiss Axio Imager M2 upright microscope equipped with an ApoTome.2. Immunohistochemical analysis was performed using Fiji (NIH) by an observer blind to genotype and experimental conditions.

### Microglia activation assessment

Microglia morphology, Iba1 content, and cytokine gene expression was determined following the initiation of the isoflurane exposure. For immunohistochemistry, brains were isolated and processed as described above 24 hours following the initiation of isoflurane exposure. Iba1 containing cells were labeled by antibody staining with 1:500 anti-Iba1 (Wako #019-19741) and subsequently 1:1000 anti-rabbit Alexa 568 and 1:5000 Hoechst 33342. Sections were imaged using a Zeiss Axio Imager M2 upright microscope equipped with an ApoTome.2 at low (10x) magnification. High magnification images were collected as a 0.5 μm z-stack with a Nikon A1R confocal microscope with an optical magnification of 60x and digital magnification of 3.5x and images were processed as a projection through the stack using Fiji.

Cortical Iba1 content was determined by western blot 24 hours following the initiation of isoflurane exposure. Cortical hemispheres including the hippocampus were surgically isolated and snap frozen in liquid nitrogen and stored at −80 °C. Tissue was lysed in 2 ml of 20 mM Tris HCl pH 7.5, 150 mM NaCl, 1 mM EDTA, 1% Triton X-100 and 1xHALT Protease and Phosphatase Inhibitor Cocktail (ThermoFisher) per hemisphere and processed with a dounce homogenizer. Insoluble material was removed by centrifugation (8000 xg for 15 minutes) and discarded. Protein concentration was determined with a Pierce BCA Protein Assay Kit (ThermoFisher) and 5 μg protein/sample were separated on 15% SDS-PAGE gels. Protein samples were transferred to PVDF membranes (ThermoFisher) and blocked in TBS Odyssey Blocking Buffer (Li-Cor). Membranes were probed with 1:2000 anti-Erk2 (Santa Cruz Biotechnology sc-1647), 1:1000 anti-Bax (CST D3R2M), and 1:500 anti-Iba1 (Wako #019-19741) followed by species-specific anti-IgG IR800CW or IR680CW secondary antibodies (Li-Cor). Samples were imaged using an Odyssey CLx system (Li-Cor). Band density quantification was performed using Fiji.

Markers of microglia activation and microglia-derived inflammatory cytokine expression were assessed by RT-PCR 12 hours after initiation of isoflurane exposure. Cortical hemispheres including the hippocampus were surgically isolated and RNA was isolated using an RNeasy Mini Kit (Qiagen). RNA concentration was determined using a NanoDrop (ThermoFisher), and 4 μg RNA was used for cDNA synthesis with a SuperScript III First-Strand Synthesis System using random hexamer primers (ThermoFisher). Template cDNA was diluted 1:5 in nuclease free H_2_O and stored at −20 °C until PCR which was performed within 7 days. Uniplex qPCR was preformed using TaqMan Fast Advanced Master Mix and FAM labeled primers (ThermoFisher) with a ViiA 7 System (Applied BioSystems). The UBE2D2 gene transcript was used as the internal reference, as it has previously been demonstrated to be relatively resistant to expression alteration in neurotoxic states(23).

### GABAergic interneuron quantification and selective protection

PND 7 mice from *Gad2*^*IRES-Cre/+*^;*Bax*^*Flox*^; *R26*^*TdTom/TdTom*^ line were used to determine the relative cortical GABAergic interneuron population size by FACS. Cortical hemispheres including the hippocampi were isolated in Hank’s Buffered Salt Solution (HBSS) at 4°C and the meninges were removed. Hemispheres were then minced to ∼1 mm^3^ pieces and digested in 1ml of HBSS, 10 units papain (Roche), 5 mM L-cysteine, and 50 units DNase (Promega) for 15 minutes at 37 °C with agitation. Digestion was halted by the addition of 100 μl fetal bovine serum (ThermoFisher). Neurons were dissociated by trituration, and the cell suspension was then filtered through a 40 μm cell strainer. The cell suspension was then floated on top of 5 ml 20% Percoll (Sigma) in HBSS and centrifuged (800 xg for 5 minutes). The supernatant was discarded, and the pellet resuspended in 0.5 ml HBSS. TdTomato positive and negative neuron populations were counted and sorted by RFP fluorescence, forward scatter and side scatter gates using a Becton Dickinson InFlux cell sorter (OHSU Flow Cytometry Core). A sample of the FACS input cell suspension as well as the sorted cells were lysed with 100 μl/100K cells of the lysis buffer described above and subjected to 10% SDS-PAGE and western blot with 1:500 anti-RFP (Rockland 8E5.G7) and 1:1000 anti-Bax antibodies as described above.

To evaluate relative vulnerability of the GABAergic interneuron population versus the non-GABAergic interneuron population following exposure to isoflurane by immunohistochemistry, the *Gad2*^*IRES-Cre/+*^;*Bax*^*Flox/Flox*^; *R26*^*TdTom/TdTom*^ and *Gad2*^*IRES-Cre/+*^;*Bax*^*Flox/+*^; *R26*^*TdTom/TdTom*^ animals were exposed to isoflurane or control conditions as described above. Two hours after conclusion of the exposure, animals were euthanized and brains sections prepared as described above. Sections were stained with 1:500 anti-RFP and 1:1000 anti-cleaved caspase 3. Cleaved caspase 3 positive neurons in cortical layers II/III were counted and the proportion of GABAergic neurons (TdTomato positive) determined by an investigator blinded to the experimental conditions.

### Seizure susceptibility assay

To investigate whether neonatal exposure to anesthesia influenced seizure susceptibility, we used a flurothyl exposure paradigm as previously described(24, 25). *Gad2*^*IRES-Cre/+*^;*Bax*^*Flox/Flox*^; *R26*^*TdTom/TdTom*^ and *Gad2*^*IRES-Cre/+*^;*Bax*^*Flox/+*^; *R26*^*TdTom/TdTom*^ animals were exposed to isoflurane or control conditions on PND7 as described above, then returned to their home cages and allowed to age until undergoing seizure susceptibility testing during post natal week (PNW) 7-8. Briefly, animals were placed in an enclosed chamber with a physically isolated vaporization chamber. Liquid 10% Bis (2,2,2-Trifluoroethyl) Ether in 85% EtOH/5% H2O solution was delivered to the vaporization chamber at a rate of 100ul/min. Time to the onset of the first myoclonic jerk and generalized tonic-clonic seizure (TCS) as evidenced by full body convulsant movements with loss of postural control was recorded by an observer blind to the experimental conditions. Verification of assay efficacy across genotype and isoflurane treatment conditions was performed through repeated daily exposures for 7 days, which was found to result in similar levels of kindling with reduced latency to TCS following repeated exposures (data not shown). Immediately following onset of TCS the animals were removed from the exposure chamber, resulting in spontaneous seizure cessation within 5 seconds. Animals recovered to their baseline activity status over 5 minutes and were then returned to their home cages.

### Behavioral Assays

The long-term consequences on learning and memory following early life exposure to isoflurane were assessed. Animals were exposed to isoflurane on PND 7 as described above and aged to PNW 10-11, at which time they underwent directed behavioral tests. Hippocampal-dependent visual-spatial learning and memory was assessed by Morris-Water-Maze (MWM) on PNW 10. Hippocampal-and amygdala-dependent contextual fear memory formation and amygdala-dependent cued fear memory were assessed on PNW 11.

MWM testing was performed generally as previously described(26). Mice were placed in a large water bath (122cm wide; 20 °C ± 1) surrounded by prominent visual cues and were removed upon locating the hidden platform (submerged 1cm under opaque water). The time taken to locate a hidden platform, the escape latency, was recorded. Mice that did not locate the platform within 60 seconds were gently guided to the platform and allowed to remain on it for 3 seconds before being removed. Each mouse received 4 trials per session with 10 minutes between each trial. Two sessions separated by 1 hour were conducted per day. After seven sessions, mice underwent a probe trial in which the hidden platform was removed. Time spent in each quadrant of the water bath was recorded. In addition, the cumulative distance the mice swam in search of the hidden platform was recorded.

Conditioned fear testing was used to assess fear-associated memory formation. On the first day of conditioned fear training, mice were placed in a novel fear conditioning chamber for 2 minutes and allowed to explore while baseline freezing time was recorded. The mice were then exposed to a 30 second tone (80dB), which was immediately followed by a 0.4mA foot shock for 2 seconds. Two minutes later, the tone-shock pairing was delivered again. Ten seconds later, the mice were removed from the chamber and returned to their home cages. The following day, mice were placed into the same fear conditioning chamber and their mobility was recorded for 3 minutes in order to assess freezing behavior (cessation of all movement except for respiration). No tones or shocks were delivered. One hour later mice were placed in a novel chamber and the same conditioned fear tone was delivered 3 minutes later. Freezing behavior was recorded during the three minutes prior to exposure to the fear tone and for the three minute period following exposure to the tone.

### Statistical analysis

Simultaneous comparison of neuron death among the genotypes and conditions tested was preformed by two-tailed Dunnett’s test using the statistical software R. Gene expression was quantified using the comparative C_T_ method in QuantStudio 6 and expressed and mean ± 95% confidence interval. A three way repeated measures ANOVA was used to determine potential group differences in the learning acquisition of the water maze test with sessions as the within subject factors and genotype and isoflurane exposure as the between subject factors. A one way ANOVA was preformed for the MWM probe trial to assess within group differences between the target quadrant versus the non-target quadrants with the statistical software SPSS. Intergroup differences in the cumulative distance swam away from the center of the target platform was assessed by two-way t-test using R.

## Results

### Bax is necessary for neuronal death following neonatal exposure to anesthesia

Neuronal death associated with ethanol exposure in early life occurs primarily via apoptosis, and is blocked by genetic deletion of Bax(10). We therefore sought to test the hypothesis that neonatal exposure to isoflurane induces cell death through apoptosis. We developed an exposure paradigm that partially titrates isoflurane delivery to a level of 1 MAC over a 6 hour period (Fig 1A). Animals tolerated this exposure well, with a low mortality rate and rapid recovery to baseline activity following completion of the exposure period. Two hours following completion of the exposure, control and isoflurane treated animals were indistinguishable based on appearance and activity level. At this point, animals were euthanized, and brains were assessed by immunohistochemistry using cleaved caspase-3 as a marker of apoptotic cell death (Fig 1B). We found that in wild-type animals exposed to isoflurane treatment, cleaved caspase-3 containing neurons were readily detectable throughout the brain. Consistent with previous descriptions, the distribution of cleaved caspase-3 neurons was higher in the superficial cortex and in layer V(9, 27). Assessment of cell death in wild-type animals using an unbiased marker of dead and degenerating neurons, FluoroJade C, revealed a pattern of neuronal death similar to that seen with cleaved caspase-3 (Fig 1C). The number of apoptotic cells was quantified by counting the number of cleaved caspase-3 positive cells with neuronal morphology within a region of interest that included the cortex and hippocampus (Fig 1D). This analysis revealed that isoflurane exposure results in a 5-fold increase in cleaved caspase-3 positive neurons compared to control treated wild-type animals (Fig 1E).

**Fig 1.**
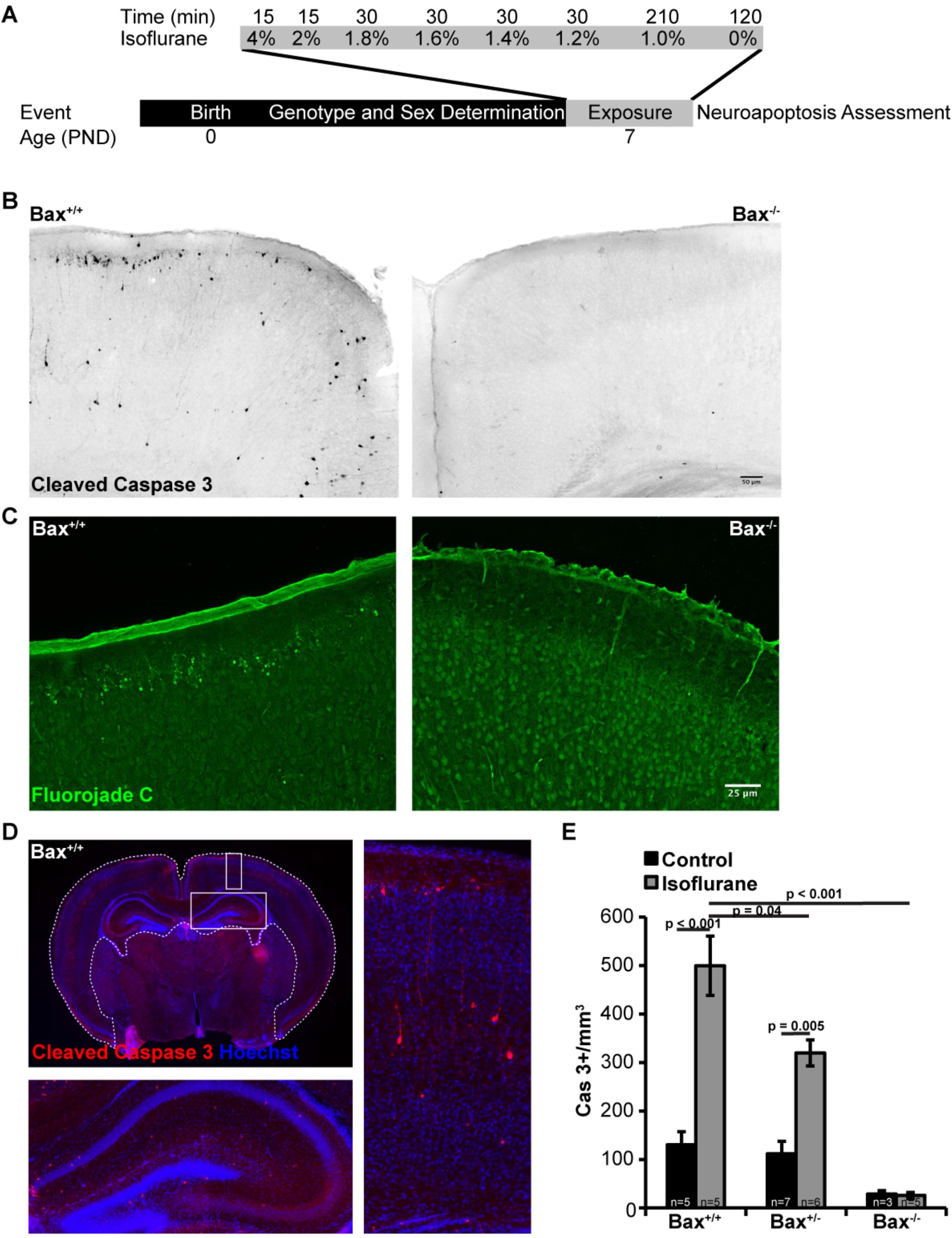
Neuronal death occurs in neonatal animals exposed to isoflurane. A) Diagram of the experimental isoflurane exposure paradigm. On PND 7 mice were exposed to isoflurane titrated to a level of 1 MAC over the duration of the exposure. The time mice were exposed to each concentration of isoflurane is shown. B) Coronal sections approximate to Paxinos plate P6 #30 from Bax^+/+^ and Bax^-/-^ animals exposed to isoflurane and probed for cleaved caspase-3. C) Coronal sections similar to (B), but stained with Fluorojade C reveals dye accumulation in neurons in the exposed Bax^+/+^ animals, but not in Bax^-/-^ animals. D) Representative brain section labeled for cleaved caspase-3 indicating the region of interest (dashed line) where cleaved caspase-3 positive neuron profiles was counted. Magnifications of boxed regions are shown below (hippocampus) and to the right (cortex). E) Quantification of cleaved caspase-3 neurons in the cortex and hippocampus.

In contrast to wild-type mice, *Bax* deficiency afforded protection from apoptotic cell death following exposure to anesthesia in a gene-dose dependent manner (Fig 1B-E). *Bax*^*+/-*^ mice had significantly fewer cleaved caspase-3 positive neurons following exposure to isoflurane than wild-type mice, (Fig. 1E). Constitutive deletion of *Bax* resulted in a nearly complete elimination of cleaved caspase-3 positive neurons following exposure to isoflurane. Cleaved caspase-3 staining was also lower in untreated *Bax*^*-/-*^ animals compared to untreated wild-type and *Bax*^*+/-*^ animals, reflecting inhibition of normal developmental apoptotic cellular pruning. Taken together, these results show that neuronal death following neonatal exposure to isoflurane occurs via Bax-dependent apoptosis.

### Neuroinflammation occurs as a consequence of Bax-mediated neuronal apoptosis

Neuroinflammation has been described following exposure to volatile anesthetic exposure in the neonatal period, and it has been proposed that inflammation itself contributes to cognitive deficits later in life(15). We therefore tested whether neuroinflammation occurs as a consequence on neuronal death, or is an independent process. We found that following exposure to isoflurane, Iba1^+^ microglia in the hippocampus (Fig 2A) and cortex (Fig 2B) of wild-type mice undergo a clear change in morphology. Microglia processes appear to retract around the soma, changing from a ramified morphology to an ameboid morphology consistent with their activation(28). In contrast, in *Bax*^*-/-*^ animals, in which neuronal apoptosis is blocked, microglia retain their ramified morphology following exposure to isoflurane. In contrast to the change in microglial morphology, the levels of Iba1 protein does not change in wild-type or *Bax*^*-/-*^ mice following exposure to isoflurane (Fig 2C).

**Fig 2.**
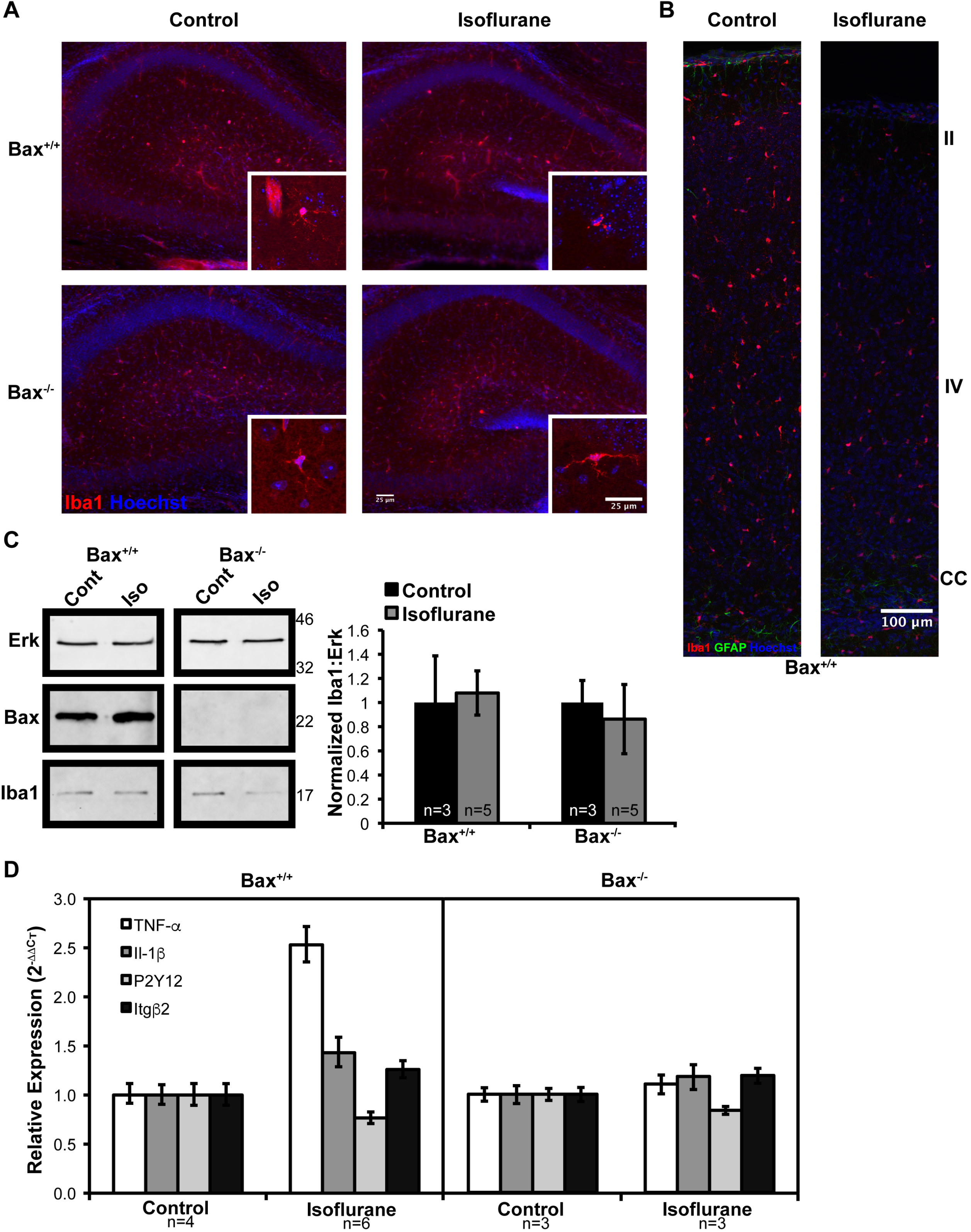
Neuroinflammation following neonatal exposure to isoflurane is a consequence of neuronal apoptosis. A) Low magnification images of hippocampal microglia and high magnification images taken in the molecular layer in the CA1 region (inset). B) Microglia imaged throughout the cortex. Approximate position of cortical layers II, IV and the corpus callosum (CC) are shown. C) Western blot analysis of Erk, Bax and Iba1 content in cortical lysate and quantification by band densitometry. D) qPCR analysis of the relative expression of microglia-derived pro-inflammatory cytokines and microglial activation genes, mean ± 95% confidence intervals are shown.

We also assessed microglial activation by quantifying the expression of the microglia-derived pro-inflammatory cytokines *TNFα* and *IL-1β*, as well as *CR3/MAC-1 β2-integrin subunit* (*Itgβ2*) and *P2Y12*, which are up-and down-regulated, respectively, during microglia activation. As expected, the expression of *TNFα, IL-1β*, and *Itgβ2* are all increased in wild-type mice following exposure to isoflurane, while *P2Y12* expression is decreased (Fig 2D). In contrast, the expression profile of microglia-activation associated genes is attenuated in *Bax*^*-/-*^ animals following isoflurane exposure (Fig 2D). These results demonstrate that microglial activation occurs downstream of Bax-mediated neuron apoptosis following neonatal exposure to anesthesia.

### GABAergic neurons are vulnerable to isoflurane-induced death

GABAergic neurons represent ∼15% of the total cortical neuron population in adult rodents. The final population of GABAergic neurons is regulated by a wave of Bax-dependent apoptosis during the early postnatal period, which overlaps with the period of vulnerability to volatile anesthetic toxicity(16, 29). Previous studies have suggested that cortical GABAergic interneurons may be overrepresented among dead cells following exposure to anesthesia(9, 30). This raises the possibility that an increase in GABAergic neuron death following exposure to anesthesia occurs due to an amplification of the normal wave of developmental apoptosis.

To test this, we first quantified the proportion of GABAergic neurons within the cleaved caspase-3 positive population following exposure to isoflurane. To identify GABergic neurons, we used a genetic approach by generating a conditional knockout/reporter line, *Gad2-IRES*^*Cre*^;*Bax*^*Flox*^;*R26*^*TdTom/TdTom*^. At PND7, the relative proportion of tdTomato+ GABAergic interneurons was ∼16-17% of the cortical neuron population based on FACS (Fig 3A). Consistent with previous reports, deletion of *Bax* did not affect overall interneuron numbers at P7, as the majority of developmental GABAergic neuron apoptosis occurs between P7-P11 (16). We observed cleaved caspase-3 staining in both GABAergic (tdTomato+) and non-GABAergic (tdTomato-) neurons in control *Gad2-IRES*^*Cre*^;*Bax*^*Flox/+*^;*R26*^*TdTom/TdTom*^ mice (Fig 3B). Quantification revealed that ∼30% of the cleaved caspase-3 neurons were tdTomato+ GABAergic neurons (Fig 3C). Therefore, GABAergic neurons were overrepresented approximately two-fold in the population of neurons undergoing apoptosis following exposure to anesthesia. Conditional deletion of *Bax* selectively from interneurons (*Gad2-IRES*^*Cre*^;*Bax*^*Flox/Flox*^;*R26*^*TdTom/TdTom*^) resulted in a 50% reduction in the proportion of GABAergic neurons undergoing apoptosis to ∼15%. Based our results using constitutive *Bax*^*-/-*^ mice (Figure 1), we had anticipated complete protection of interneurons from anesthesia-associated death in these mice. However, when we assessed the degree of *Bax* deletion in *Gad2-IRES*^*Cre*^;*Bax*^*Flox/Flox*^;*R26*^*TdTom/TdTom*^ animals, we found that Bax protein in interneurons was reduced, but not eliminated (Fig 3D). This suggests that despite the *Gad2* promoter driving recombination at ∼E19(31), there was still residual Bax protein at PND7, possibly due to slow protein turnover. Regardless, even incomplete elimination of Bax protein resulted in significant protection from isoflurane-induced apoptosis in GABAergic neurons.

**Fig 3.**
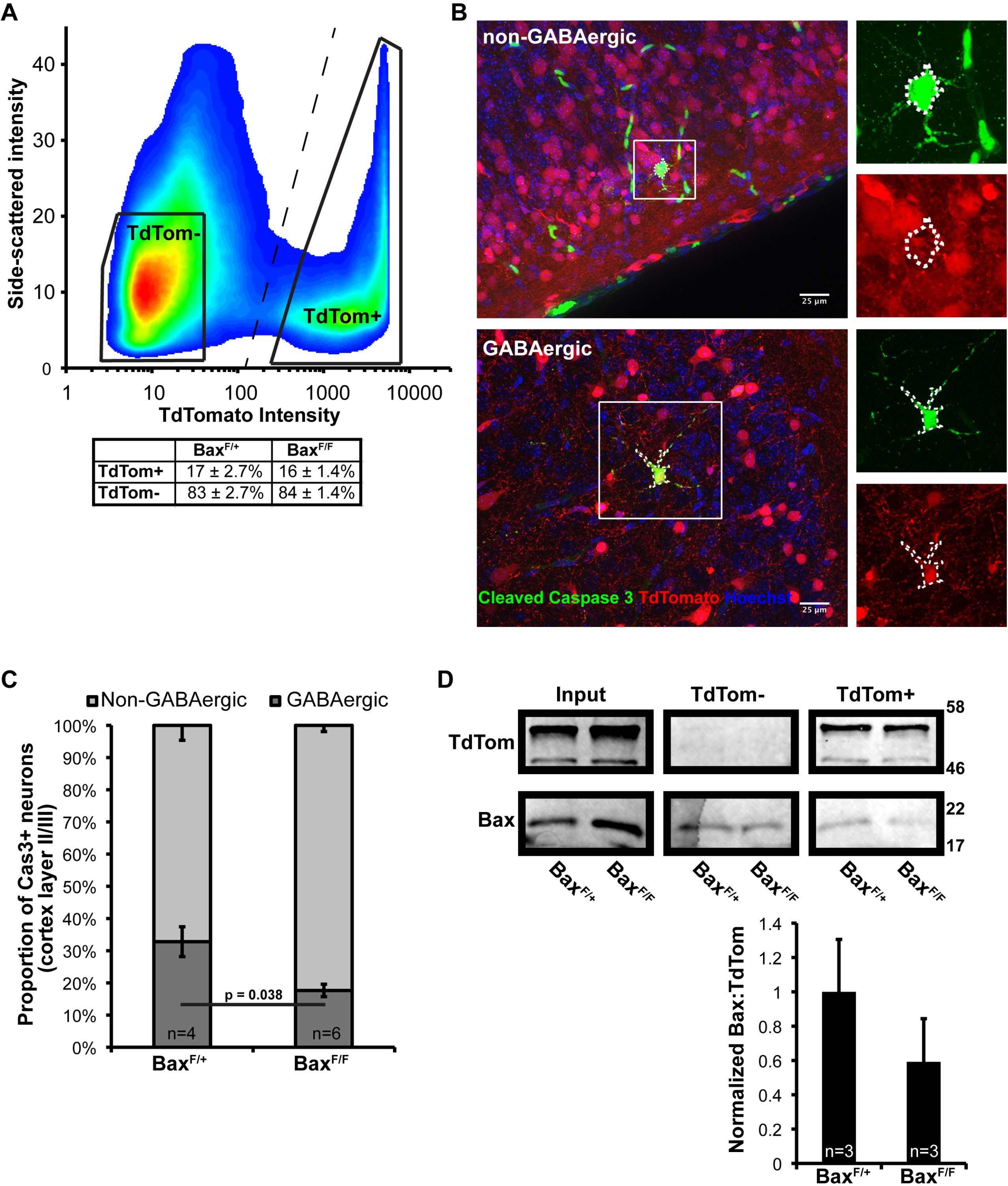
GABAergic interneurons show increased vulnerability to isoflurane-induced apoptosis. A) Cortical interneurons from the Gad2-IRES^Cre^;Bax^Flox^;^TdTom/TdTom^ were counted by FACS based on TdTomato fluorescence and side scatter. Demarcation of the GABAergic and nonGABAergic neuron populations for counting is shown as the dashed line. Solid trapazoids are representative of the collection gates used for sorting the cell populations used for subsequent western blot analysis in (D). Three animals per genotype from three separate litters were used for this analysis. B) Representative images of cleaved caspase-3 positive non-GABAergic (top) and GABAergic (bottom) neurons following exposure to isoflurane. C) Quantification of the relative proportion of non-GABAergic and GABAergic neurons in the total cleaved caspase-3 positive population in cortical layers II/III. D) Western blot analysis of input and the isolated populations following FACS with quantification of the relative Bax levels in the TdTomato positive population by band densitometry.

### Selective protection of GABAergic neurons alters seizure susceptibility

Prior work suggests that exposure to volatile anesthetics may result in lasting alteration of seizure susceptibility. First, volatile anesthetics cause epileptic discharge-like activity when delivered in sub-burst suppression doses(32, 33). Furthermore, over activation of microglia during early development is known to enhance epileptogenicity of neurotoxic insults(34). Finally, disruption of the excitatory:inhibitory (E:I) ratio through loss of GABAergic interneurons correlates with epilepsy severity(35). Given our observations that isoflurane exposure activates microglia and that GABAergic neurons are overrepresented among cleaved caspase-3 positive neurons, we tested whether the mice exposed to isoflurane as neonates showed altered seizure susceptibility as adults. We also asked whether selectively reducing inhibitory neuron death in *Gad2-IRES*^*Cre/+*^;*Bax*^*Flox/Flox*^ mice offered protection from seizure susceptibility.

We induced seizures with exposure to Bis-(2,2,2-Trifluoroethyl)-Ether (Flurothyl) due to its ease of administration, demonstrated consistency, and rapid recovery from seizure following cessation of exposure(24, 25). Latency to the onset of the first myoclonic jerk following exposure to Flurothyl was not statistically different between control and isoflurane exposed groups in either *Gad2-IRES*^*Cre/+*^;*Bax*^*Flox/+*^ or *Gad2-IRES*^*Cre/+*^;*Bax*^*Flox/Flox*^ (Fig 4A). Similar to the results seen with latency to the first myoclonic jerk, control *Gad2-IRES*^*Cre/+*^;*Bax*^*Flox/+*^ mice exposed to isofluane did not display any difference in latency to tonic-clonic seizure (TCS). Surprisingly, isoflurane exposed *Gad2-IRES*^*Cre/+*^;*Bax*^*Flox/Flox*^ mice demonstrated consistent protection from TCS induction (Fig 4B). Taken together, these results suggest that the increased neuron apoptosis and neuroinflammation observed following early life exposure to isoflurane does not alter susceptibility to seizures later in life.

**Fig 4.**
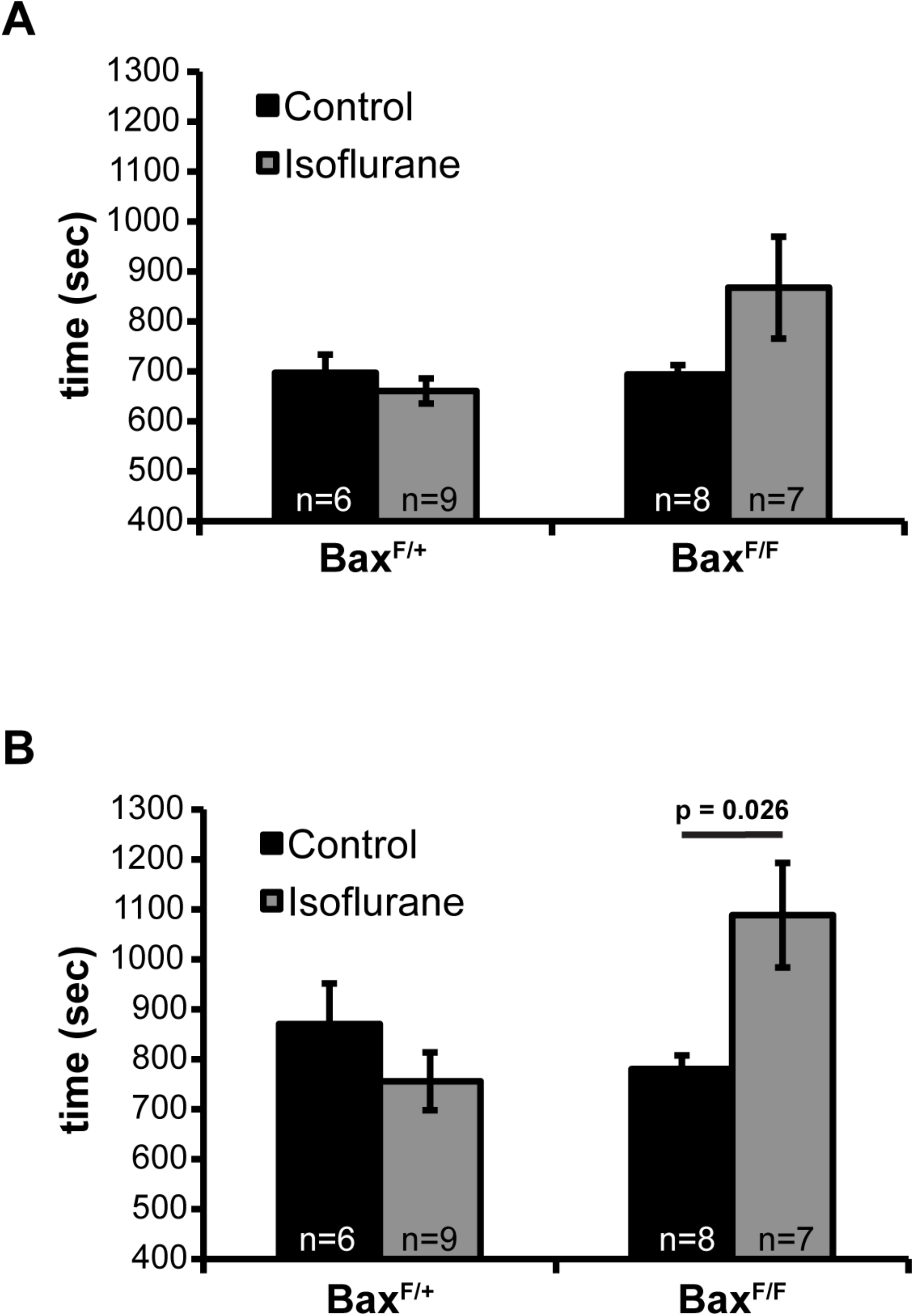
Early life exposure to isoflurane does not affect seizure susceptibility. A) Latency to the onset of the first myoclonic jerk following the initiation of the exposure to Fluothyl. B) Latency to the onset of TCS with loss of postural control.

### Assessment of learning and memory following isoflurane exposure in early life

We found that exposure to isoflurane early in life induced neuronal apoptosis and neuroinflammation, which could be blocked by constitutive deletion of *Bax*. Therefore, we tested whether neonatal exposure to isoflurane causes cognitive defects later in life, and whether blocking neuronal apoptosis in this context provides protection. We used the Morris Water Maze (MWM) test to assess deficits in hippocampal-dependent visual-spatial memory formation following early life exposure to anesthesia(1). There was no effect on the escape latency during the training period when comparing controls and mice exposed to isoflurane in the *Bax*^*+/+*^, *Bax*^*+/-*^, and *Bax*^*-/-*^ groups (Fig 5A). Unexpectedly, unexposed *Bax*^*-/-*^ animals showed an increase in escape latency compared to unexposed *Bax*^*+/+*^ animals through the first six training sessions. A probe trial was performed 24 hours following the training sessions to assess memory retention and recall (Fig 5B). All genetic and treatment conditions displayed place preference for the quadrant formerly containing the hidden platform (the target quadrant). We assessed potential group differences in the memory retention and recall process by evaluating cumulative distance the animal swam from the center point of the hidden platform (Fig 5C). There was no difference between control and isoflurane exposed animals within each genotype group. However, a significant difference between the *Bax*^*+/+*^ and *Bax*^*-/-*^ groups was observed, with *Bax*^*-/-*^ animals swimming a greater cumulative distance.

**Fig 5.**
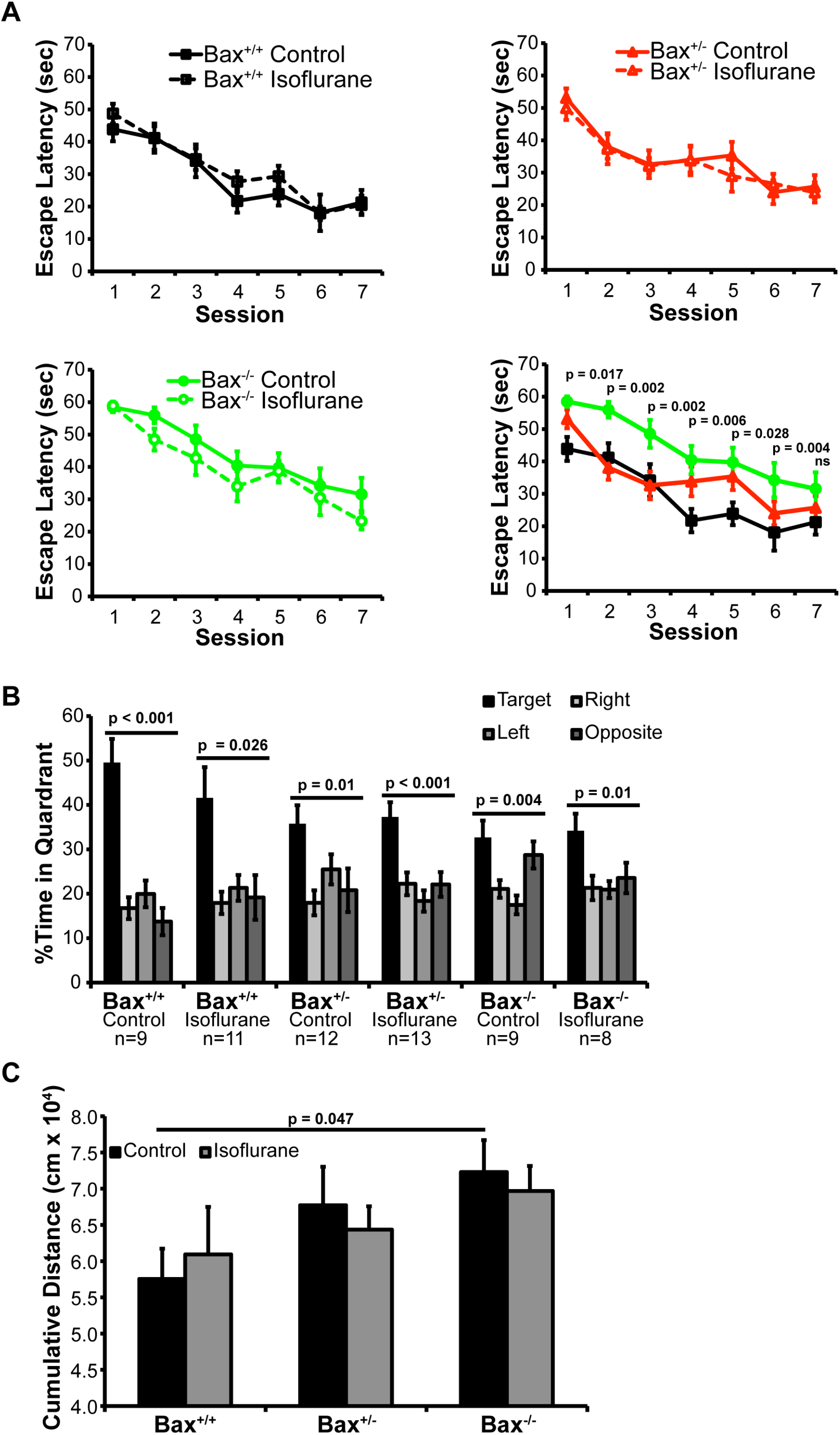
Hippocampal-dependent visual spatial memory is unaffected by early life exposure to isoflurane. A) Learning curves on repeated training sessions in the Morris Water Maze, group sizes are as indicated in (B). B) Place preference assessed as percent time spent in each quadrant of the maze over the 60 seconds of the probe trial in which the escape platform was removed and the visual cues remained unchanged. C) Cumulative distance the animal swam from where the platform was originally located.

In addition to the MWM, fear conditioning assays have been used extensively to evaluate cognitive deficits in rodents exposed to volatile anesthetics in early life(1). We used a standard training regime and tested control and isoflurane-exposed Bax^+/-^, Bax^+/-^ and Bax^-/-^ mice 24 hours later for contextual and cued fear behavior. During the training period, we observed no differences in freezing time or mobility among the animals for the first two minutes the subjects were in the novel fear conditioning chamber prior to exposure to noxious auditory or electrical stimuli (Fig 6A). One day following training, we assessed contextual fear memory and observed no difference between control and isoflurane exposed animals within each genotype group (Fig 6B). However, similar to the MWM test, *Bax*^*-/-*^ mice from the control group exhibited a weaker memory, characterized by less freezing time. Cued fear memory displayed a similar pattern (Fig 6C). Although the *Bax*^*-/-*^ mice did not show a significant increase in freezing upon re-exposure to the tone that was previously associated with the shock, there were no significant interactions between genotype and treatment to suggest that this was independent of its genotype deficit.

**Fig 6.**
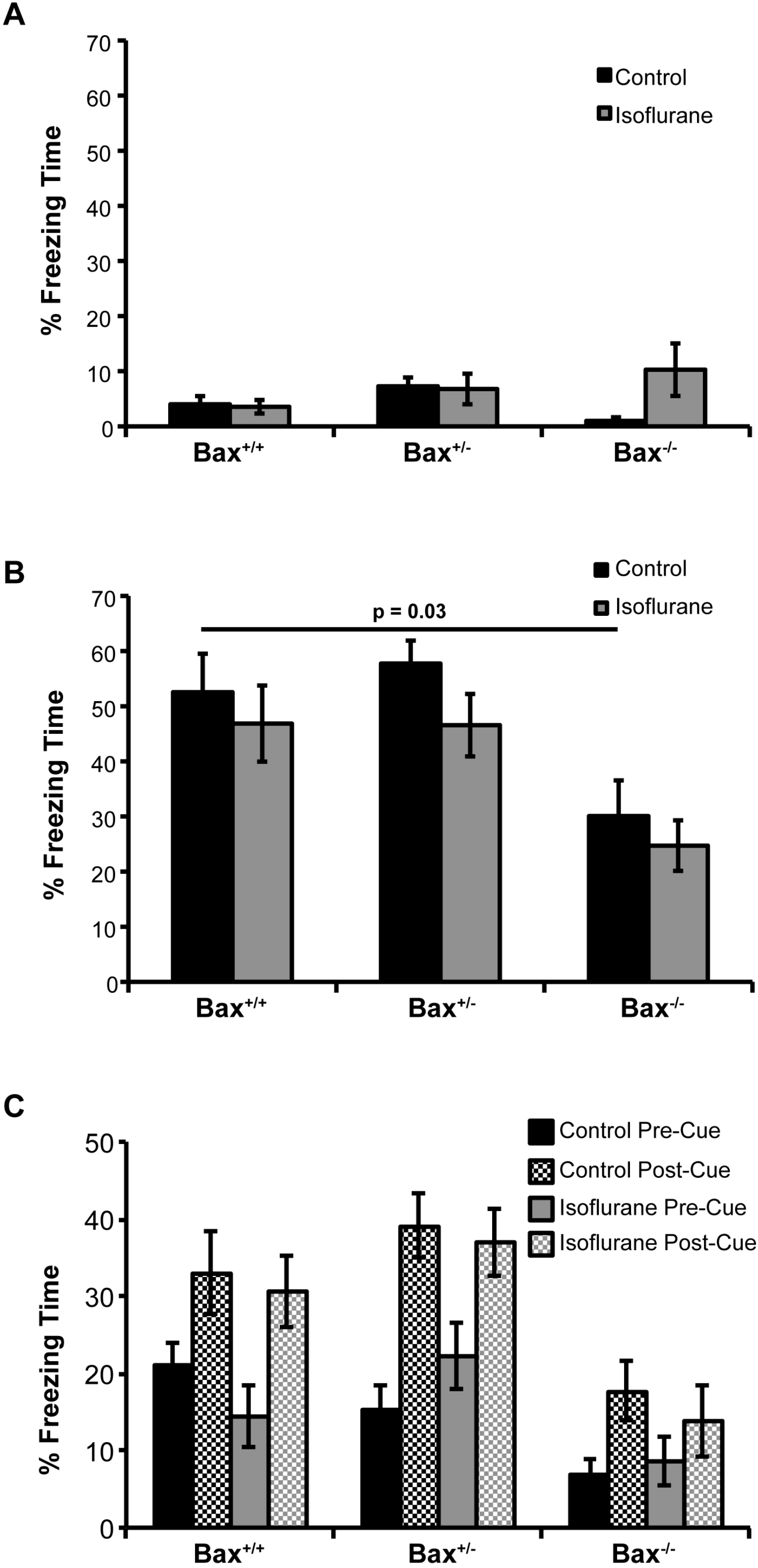
Hippocampal/Amygdala-dependent fear conditioning is unaffected by early life exposure to isoflurane. A) Baseline freezing time assessed during the first exposure to the fear conditioning chamber prior to exposure to noxious stimuli. B) 24 hours after training, mice were returned to the fear conditioning chamber and freezing time over a 3 minute observation period was assessed. Comparison between the Bax^+/+^ and Bax^-/-^ groups that were significant are indicated. C) One hour after contextual fear assessment, mice were placed in a novel chamber and exposed to the auditory tone heard during training. Freezing time prior to exposure and following the tone was assessed.

Collectively, these results show that early life exposure to isoflurane did not result in deficits in hippocampal-dependent visual spatial memory formation or hippocampal/amygdala-dependent fear conditioning. Interestingly, we did observe defects in *Bax*^*-/-*^ mice in the control groups, suggesting that interfering with normal developmental apoptosis resulted in memory-related cognitive deficits.

## Discussion

Previous studies have indicated that in addition to neuronal death, exposure to anesthetic agents in the neonatal period results in disrupted synapse architecture, altered mitochondrial morphology and increased axonal pruning(36-38). We attempted to delineate the contribution of neuronal apoptosis from other potentially injurious effects associated with exposure to volatile anesthetics in the neonatal period using *Bax* knockout mice. We found that Bax is essential for isoflurane-induced neuronal death, as *Bax*^*-/-*^ mice displayed no evidence of apoptotic or dead/degenerating neurons following exposure to isoflurane at PND7.

Neuroinflammation has been described in adult models of anesthesia exposure, but has not been assessed in neonatal models(39). We therefore investigated whether neuroinflammation occurs following early life exposure to isoflurane, and whether it occurs independent of neuronal death using *Bax* knockout mice. In control mice, there were clear changes in microglial morphology consistent with their activation immediately following exposure to isoflurane. This was accompanied by changes in pro-inflammatory cytokines associated with microglial activation. This inflammatory response was dependent on neuronal apoptosis, as it was attenuated in *Bax* knockout mice. This suggests that during the neonatal period, volatile anesthetics are not a strong proinflammatory stimuli themselves, but rather neuronal apoptosis is the primary stimuli which induces microglial activation. This is similar to the injury and response pattern observed with early life exposure to ethanol(14).

Bax-dependent cellular pruning is a normal part of neurological development, and GABAergic neurons undergo extensive apoptotic cellular pruning during the first two weeks of post-natal life(16). Cortical GABAergic interneurons are also thought to be particularly vulnerable to volatile anesthetic associated death in the neonatal period(30). Indeed, we found that GABAergic interneurons are overrepresented within the apoptotic population following exposure to isoflurane. We predicted that excessive interneuron apoptosis could potentially lead to disruptions in E:I balance and increased susceptibility to seizures. Supporting this hypothesis, epileptogenic-like EEG has been observed as a consequence of early life exposure to volatile anesthetics(40, 41). However, we found no decrease in seizure threshold associated with early life exposure to volatile anesthetics in control genotype mice. One potential explanation for this unexpected result is that the acute increase in GABAergic neuronal apoptosis immediately following exposure to isoflurane at P7 is balanced by a reduction of GABAergic apoptosis in the later stages of normal development (P10-P14), resulting in an overall preservation of GABAergic neuron numbers. Alternatively, overall numbers of GABAergic neurons may be decreased following early life exposure to isoflurane, with the remaining GABAergic neurons scaling their inhibitory inputs to maintain a normal E:I balance. We should also note that our study differed from previous work in the anesthetic used (sevoflurane vs isoflurane), which may account for the different results(41). Surprisingly, we did find that *Gad2-IRES*^*Cre/+*^;*Bax*^*Flox/Flox*^ mice had increased seizure threshold in isoflurane exposed mice, suggesting that selective protection of GABAergic neurons from volatile anesthetic associated death results in alterations of the global E:I ratio.

We also investigated whether neuronal apoptosis and neuroinflammation following early life exposure to volatile anesthetics had lasting consequences on cognitive function later in life. We found no significant differences between controls and isoflurane-exposed wild-type animals in the MWM or fear conditioning assays, suggesting that early life exposure does not affect cognitive function. To our surprise, we found that *Bax* deficiency itself led to deficits in multiple aspects of cognition, independent of volatile anesthetic exposure. Previous studies have described *Bax* knockout mice as having persistently prolonged escape latency in MWM training sessions when visual spatial learning was assessed(42, 43). The deficits we observed suggest that normal Bax-mediated developmental apoptosis in early life is critical for cognitive development. Cellular pruning, synapse remodeling, and axon pruning are all mediated by Bax-dependent caspase activation, and disruption of any or all of these processes could give rise to negative effects on cognitive development(16, 44, 45). We cannot exclude the possibility that the lack of cognitive deficits in wild-type animals exposed to isoflurane treatment in early life is due to specific aspects of the study design. However, if the neuronal apoptosis and neuroinflammation attributable to volatile anesthetic exposure in early life does contribute to cognitive defects later in life, it is relatively mild when compared to the effect that blocking normal developmental caspase activation with *Bax* deficiency has on cognitive development.

In conclusion we have demonstrated that Bax is necessary for neuronal death associated with early life exposure to volatile anesthetics. The neuroinflammation seen following volatile anesthetic exposure in the neonatal period likely arises as a secondary consequence of Bax-mediated neuronal apoptosis rather than being an independent response to volatile anesthetic exposure. And finally, GABAergic interneurons are overrepresented among the dead neurons, suggesting they are more susceptible to the pro-apoptotic effects of volatile anesthesia. Due to the cognitive deficits attributable to *Bax* deficiency alone, we were unable to conclusively determine whether blocking apoptosis and neuroinflammation following volatile anesthetic exposure provided any benefit with respect to cognitive function. Establishing a transient protected state through the use of Bax- or caspase-specific small molecule inhibitors may prove to be a viable alternative approach for investigating lasting consequences of early life exposure to volatile anesthetics. Further investigation along these lines will be necessary to delineate the contribution of neuronal death from other disruptions in neuronal function to the injury arising from early life exposure to volatile anesthetics.

## Acknowledgements

The authors would like to thank Kylee Rosette for technical assistance with mouse colony maintenance and Mike Jacobson for technical assistance with behavioral assays. This work was supported by FAER Research Fellowship Grant (RFG-02-15-17) to AMS and NIH Grant (R01 NS091027) to KMW.

## References

1. Walters JL, Paule MG. Review of preclinical studies on pediatric general anesthesia-induced developmental neurotoxicity. Neurotoxicol Teratol. 2017;60:2–23.

2. Yon JH, Daniel-Johnson J, Carter LB, Jevtovic-Todorovic V. Anesthesia induces neuronal cell death in the developing rat brain via the intrinsic and extrinsic apoptotic pathways. Neuroscience. 2005;135(3):815–27.

3. Schenning KJ, Noguchi KK, Martin LD, Manzella FM, Cabrera OH, Dissen GA, et al. Isoflurane exposure leads to apoptosis of neurons and oligodendrocytes in 20- and 40-day old rhesus macaques. Neurotoxicology and Teratology. 2017;60:63–8.

4. Jevtovic-Todorovic V. Exposure of Developing Brain to General AnesthesiaWhat Is the Animal Evidence? Anesthesiology. 2018;128(4):832–9.

5. Warner DO, Shi Y, Flick RP. Anesthesia and Neurodevelopment in Children: Perhaps the End of the Beginning. Anesthesiology. 2018;128(4):700–3.

6. Zhang Y, Dong Y, Wu X, Lu Y, Xu Z, Knapp A, et al. The mitochondrial pathway of anesthetic isoflurane-induced apoptosis. J Biol Chem. 2010;285(6):4025–37.

7. Li Y, Zeng M, Chen W, Liu C, Wang F, Han X, et al. Dexmedetomidine Reduces Isoflurane-Induced Neuroapoptosis Partly by Preserving PI3K/Akt Pathway in the Hippocampus of Neonatal Rats. PLOS ONE. 2014;9(4):e93639.

8. Yan H, Xu T, Zhao H, Lee KC, Wang HY, Zhang Y. Isoflurane increases neuronal cell death vulnerability by downregulating miR-214. PLoS One. 2013;8(2):e55276.

9. Brambrink AM, Evers AS, Avidan MS, Farber NB, Smith DJ, Zhang X, et al. Isoflurane-induced neuroapoptosis in the neonatal rhesus macaque brain. Anesthesiology. 2010;112(4):834–41.

10. Young C, Klocke BJ, Tenkova T, Choi J, Labruyere J, Qin YQ, et al. Ethanol-induced neuronal apoptosis in vivo requires BAX in the developing mouse brain. Cell death and differentiation. 2003;10(10):1148–55.

11. Schafer ZT, Kornbluth S. The Apoptosome: Physiological, Developmental, and Pathological Modes of Regulation. Developmental Cell. 2006;10(5):549–61.

12. Deckwerth TL, Elliott JL, Knudson CM, Johnson EM, Jr., Snider WD, Korsmeyer SJ. BAX is required for neuronal death after trophic factor deprivation and during development. Neuron. 1996;17(3):401–11.

13. Xiang H, Kinoshita Y, Knudson CM, Korsmeyer SJ, Schwartzkroin PA, Morrison RS. Bax involvement in p53-mediated neuronal cell death. J Neurosci. 1998;18(4):1363–73.

14. Ahlers KE, Karacay B, Fuller L, Bonthius DJ, Dailey ME. Transient activation of microglia following acute alcohol exposure in developing mouse neocortex is primarily driven by BAX-dependent neurodegeneration. Glia. 2015;63(10):1694–713.

15. Shen X, Dong Y, Xu Z, Wang H, Miao C, Soriano SG, et al. Selective anesthesia-induced neuroinflammation in developing mouse brain and cognitive impairment. Anesthesiology. 2013;118(3):502–15.

16. Southwell DG, Paredes MF, Galvao RP, Jones DL, Froemke RC, Sebe JY, et al. Intrinsically determined cell death of developing cortical interneurons. Nature. 2012;491(7422):109–13.

17. Lee V, Maguire J. Impact of inhibitory constraint of interneurons on neuronal excitability. Journal of neurophysiology. 2013;110(11):2520–35.

18. Loepke AW, Istaphanous GK, McAuliffe JJ, 3rd, Miles L, Hughes EA, McCann JC, et al. The effects of neonatal isoflurane exposure in mice on brain cell viability, adult behavior, learning, and memory. Anesth Analg. 2009;108(1):90–104.

19. Lee BH, Chan JT, Hazarika O, Vutskits L, Sall JW. Early exposure to volatile anesthetics impairs long-term associative learning and recognition memory. PLoS One. 2014;9(8):e105340.

20. McFarlane L, Truong V, Palmer JS, Wilhelm D. Novel PCR assay for determining the genetic sex of mice. Sex Dev. 2013;7(4):207–11.

21. Paxinos G. Atlas of the developing mouse brain at E17.5, P0 and P6. 1st ed. Amsterdam; Boston: Elsevier; 2007. xi, 353 p. p.

22. Ehara A, Ueda S. Application of Fluoro-Jade C in acute and chronic neurodegeneration models: utilities and staining differences. Acta Histochem Cytochem. 2009;42(6):171–9.

23. Rydbirk R, Folke J, Winge K, Aznar S, Pakkenberg B, Brudek T. Assessment of brain reference genes for RT-qPCR studies in neurodegenerative diseases. Sci Rep. 2016;6:37116.

24. Papandrea D, Anderson TM, Herron BJ, Ferland RJ. Dissociation of seizure traits in inbred strains of mice using the flurothyl kindling model of epileptogenesis. Exp Neurol. 2009;215(1):60–8.

25. Ferland RJ. The Repeated Flurothyl Seizure Model in Mice. Bio Protoc. 2017;7(11).

26. Morris R. Developments of a water-maze procedure for studying spatial learning in the rat. Journal of neuroscience methods. 1984;11(1):47–60.

27. Istaphanous GK, Ward CG, Nan X, Hughes EA, McCann JC, McAuliffe JJ, et al. Characterization and quantification of isoflurane-induced developmental apoptotic cell death in mouse cerebral cortex. Anesth Analg. 2013;116(4):845–54.

28. Beynon SB, Walker FR. Microglial activation in the injured and healthy brain: what are we really talking about? Practical and theoretical issues associated with the measurement of changes in microglial morphology. Neuroscience. 2012;225:162–71.

29. Meinecke DL, Peters A. GABA immunoreactive neurons in rat visual cortex. J Comp Neurol. 1987;261(3):388–404.

30. Zhou ZW, Shu Y, Li M, Guo X, Pac-Soo C, Maze M, et al. The glutaminergic, GABAergic, dopaminergic but not cholinergic neurons are susceptible to anaesthesia-induced cell death in the rat developing brain. Neuroscience. 2011;174:64–70.

31. Taniguchi H, He M, Wu P, Kim S, Paik R, Sugino K, et al. A resource of Cre driver lines for genetic targeting of GABAergic neurons in cerebral cortex. Neuron. 2011;71(6):995–1013.

32. Iijima T, Nakamura Z, Iwao Y, Sankawa H. The epileptogenic properties of the volatile anesthetics sevoflurane and isoflurane in patients with epilepsy. Anesth Analg. 2000;91(4):989–95.

33. Voss LJ, Sleigh JW, Barnard JP, Kirsch HE. The howling cortex: seizures and general anesthetic drugs. Anesth Analg. 2008;107(5):1689–703.

34. Kim I, Mlsna LM, Yoon S, Le B, Yu S, Xu D, et al. A postnatal peak in microglial development in the mouse hippocampus is correlated with heightened sensitivity to seizure triggers. Brain Behav. 2015;5(12):e00403.

35. Buckmaster PS, Abrams E, Wen X. Seizure frequency correlates with loss of dentate gyrus GABAergic neurons in a mouse model of temporal lobe epilepsy. Journal of Comparative Neurology. 2017;525(11):2592–610.

36. Boscolo A, Milanovic D, Starr JA, Sanchez V, Oklopcic A, Moy L, et al. Early exposure to general anesthesia disturbs mitochondrial fission and fusion in the developing rat brain. Anesthesiology. 2013;118(5):1086–97.

37. Amrock LG, Starner ML, Murphy KL, Baxter MG. Long-term effects of single or multiple neonatal sevoflurane exposures on rat hippocampal ultrastructure. Anesthesiology. 2015;122(1):87–95.

38. Obradovic AL, Atluri N, Dalla Massara L, Oklopcic A, Todorovic NS, Katta G, et al. Early Exposure to Ketamine Impairs Axonal Pruning in Developing Mouse Hippocampus. Mol Neurobiol. 2017.

39. Jackson WM, Gray CD, Jiang D, Schaefer ML, Connor C, Mintz CD. Molecular Mechanisms of Anesthetic Neurotoxicity: A Review of the Current Literature. J Neurosurg Anesthesiol. 2016;28(4):361–72.

40. Edwards DA, Shah HP, Cao W, Gravenstein N, Seubert CN, Martynyuk AE. Bumetanide alleviates epileptogenic and neurotoxic effects of sevoflurane in neonatal rat brain. Anesthesiology. 2010;112(3):567–75.

41. Seubert CN, Zhu W, Pavlinec C, Gravenstein N, Martynyuk AE. Developmental effects of neonatal isoflurane and sevoflurane exposure in rats. Anesthesiology. 2013;119(2):358–64.

42. Tehranian R, Rose ME, Vagni V, Pickrell AM, Griffith RP, Liu H, et al. Disruption of Bax protein prevents neuronal cell death but produces cognitive impairment in mice following traumatic brain injury. J Neurotrauma. 2008;25(7):755–67.

43. Lee JW, Kim WR, Sun W, Jung MW. Role of dentate gyrus in aligning internal spatial map to external landmark. Learn Mem. 2009;16(9):530–6.

44. Li Z, Jo J, Jia JM, Lo SC, Whitcomb DJ, Jiao S, et al. Caspase-3 activation via mitochondria is required for long-term depression and AMPA receptor internalization. Cell. 2010;141(5):859–71.

45. Simon DJ, Weimer RM, McLaughlin T, Kallop D, Stanger K, Yang J, et al. A caspase cascade regulating developmental axon degeneration. J Neurosci. 2012;32(49):17540–53.

